# YAP and TAZ differentially regulate postnatal cortical progenitor proliferation and astrocyte differentiation

**DOI:** 10.1101/2023.07.25.550567

**Authors:** Jessie Chen, Yung-Hsu Tsai, Anne Linden, John A Kessler, Chian-Yu Peng

**Affiliations:** Department of Neurology, Northwestern University’s Feinberg School of Medicine, Chicago, Illinois, 60611

**Keywords:** TAZ, BMP, YAP, integrins, astrocytes, neural stem cells

## Abstract

WW domain-containing transcription regulator 1 (TAZ) and Yes-associated protein (YAP) are transcriptional co-activators traditionally studied together as a part of the Hippo pathway and best known for their roles in stem cell proliferation and differentiation. Despite their similarities, TAZ and YAP can exert divergent cellular effects by differentially interacting with other signaling pathways that regulate stem cell maintenance or differentiation. In the developing central nervous system, In this study, we show that TAZ regulates astrocytic differentiation and maturation of postnatal neural stem and progenitor cells (NPCs), and that TAZ mediates some but not all of the effects of bone morphogenetic protein (BMP) signaling on astrocytic development. By contrast, TAZ and YAP both mediate effects on NPC fate of β1-integrin and integrin-linked kinase (ILK) signaling, and these effects are dependent on extracellular matrix (ECM) cues. These findings demonstrate that TAZ and YAP perform divergent functions in the regulation of astrocyte differentiation, where YAP regulates cell cycle states of astrocytic progenitors and TAZ regulates differentiation and maturation from astrocytic progenitors into astrocytes.

**Summary Statement:** Astrocytes are accounts for nearly half of the cells in the central nervous system, where they perform a diverse array of physiological functions. During development, astrocytes are primarily generated after neuronal differentiation in a stepwise manner from multiple glial committed progenitor subtypes. How gliogenic progenitors maintain proliferative properties versus differentiate into astrocytes is not fully understood. This work aims to elucidate how environmental signals utilizes molecularly similar intracellular components to achieve distinct developmental outcomes. In addition, many of the cell types that are involved in glial development are also present in brain tumors including glioblastoma. Knowledge on mechanisms regulating proliferation and differentiation of glial progenitors will provide insights into differences and similarities between normal and malignant cells.

## Introduction

WW domain-containing transcription regulator 1 (WWTR1 or TAZ) and Yes-associated protein (YAP) are evolutionarily conserved transcriptional co-activators found in multiple cell types. Initially identified as binding partners of Yes and 14-3-3, respectively (Sudol, 1994; Kanai et al., 2000), TAZ and YAP cannot directly bind DNA, but instead recruit and interact with other transcription factors to regulate both cell proliferation and organ size during development (Huang et al., 2005; Dong et al., 2007; Gill et al., 2018). Due to their large degree of homology, TAZ and YAP are commonly assumed to have overlapping functions. However, recent studies have indicated that they may in fact play divergent roles, depending on the biological system and timing during development(Sun et al., 2017; Plouffe et al.; 2018, Muppala et al.; 2019). Regulation of TAZ and YAP also may differ, with TAZ having additional phosphorylation sites for degradation (Huang et al., 2012). More importantly, TAZ and YAP have non-overlapping transcriptional targets that may lead to distinct physiological outcomes in response to specific environmental stimuli, such as in the context of cancer (Shreberk-Shaked et al., 2020; He et al., 2021). Whether TAZ and YAP exert distinct functions in the mammalian nervous system has not been explored.

In the developing mouse nervous system, gliogenesis in the cerebral cortex primarily takes place after embryonic neurogenesis, in a multi-step process that involves glial lineage commitment of neural progenitor cells (NPCs) in the subventricular zone (SVZ) followed by differentiation of local astrocyte precursors in the brain parenchyma. YAP has been found to promote neural stem and progenitor cells (NPCs) proliferation through action with TEA domain family transcription factors (TEADs) (Cao et al., 2008; Yao et al., 2014; Han et al., 2015). In addition to regulating NPC proliferation, YAP has also been demonstrated to promote astrocyte differentiation (Huang et al., 2016a, Huang et al., 2016b). Loss of TAZ in NPCs has been demonstrated to impair neuronal differentiation (Robledinos-Anton et al., 2020), but its role in glial fate specification and astrocyte differentiation has not been independently studied.

While TAZ and YAP are canonical effectors of Hippo signaling and are regulated by LATS phosphorylation, emerging evidence supports a role for TAZ and YAP in other signaling pathways. One candidate regulatory pathway is bone morphogenetic protein (BMP) signaling. BMP signaling is known to promotes cell cycle exit (Bond et al., 2014) and astrocyte lineage commitment in postnatal NSCs (Gross et al., 1996; Bonaguidi et al., 2005; 2008). Prior studies suggested that BMP-induced cell cycle exit of embryonic NPCs is in part due to inhibition of YAP-mediated transcriptional regulation. (Yao et al., 2014). However, YAP signaling was also reported to mediate BMP-induced astrocyte differentiation of embryonic NPCs (Huang et al., 2016b). This role of YAP in mediating BMP-induced astrocyte differentiation seemed at odds with the pro-proliferative effects of YAP described in NPCs and many types of stem cells (Kostic et al., 2019)

TAZ and YAP are characterized as mechanosensing proteins (Dupont et al., 2011; Shimizu et al., 2017) that likely also integrate mechanical cues into NPC fate decisions. There are a number of transmembrane receptors that detect the extracellular matrix and that are potential candidates for TAZ/YAP regulators, including β1-integrin. β1-integrin signaling regulates postnatal and adult NPC maintenance and suppresses NPC astrocytic lineage commitment (Leone et al., 2005; Pan et al., 2014; North et al., 2015; Brooker et al., 2016), and has been shown to modulate BMP signaling (Du, 2011, Ashe, 2016). Since components of β1-integrin signaling have also been identified as Hippo regulators in other cell systems (Serrano et al., 2013; Elbediwy et al., 2016), it is a prime candidate for regulation of TAZ and YAP in NPCs.

Although available evidence suggested that TAZ and YAP might be points of convergence for multiple signaling cues, whether TAZ and YAP differentially respond to these signals and exert distinct cellular responses in NPCs is unclear. In this study, we examined if TAZ is an effector of the BMP and β1-integrin signaling pathways in postnatal NPCs, and whether YAP and TAZ are responsible for the effects of these signaling cues on NPC proliferation and fate specification. In contrast to prior studies that demonstrated embryonic ablation of YAP impair astrocyte differentiation in NPCs (Yao et al., 2014; Huang et al., 2016a), we found that the effects of BMP signaling on postnatal astrocyte differentiation involve TAZ but not YAP. Further, while both YAP and TAZ are targets of β1-integrin/integrin-linked kinase (ILK), YAP promotes proliferation whereas TAZ promotes differentiation into glial fibrillary acidic protein (GFAP)-expressing astrocytes. These findings separate YAP and TAZ as distinct entities with unique roles in NPC proliferation and fate determination.

## Materials and Methods

### Mouse lines

Yap^tm1Hmc^;Taz^tm1Hmc^ mice were obtained from Jackson Laboratories (strain 030532). To obtain TAZ conditional knockout mice, Yap^tm1Hmc^;Taz^tm1Hmc^ mice were crossed to a C57BL/6J mouse, and progeny crossed until the YAP mutant allele was removed. Yap^dupa^ mice were obtained from Jackson Laboratories (strain 027929). Rosa26:ZsGreen (strain 007906) mice on a BL6C57 background were crossed with Yap^tm1Hmc^;Taz^tm1Hmc^ mice. Wild-type C57BL/6J mice were ordered as timed pregnancies from Jackson Laboratories. To obtain conditional knockout of YAP/TAZ in neural stem/progenitor cells, Nestin-CreERT2 (strain 016261) mice were crossed with Yap^tm1Hmc^;Taz^tm1Hmc^; Rosa26:ZsGreen mice. Postnatal day 1 mice were injected with 20 μL of tamoxifen (30 mg/ml) subcutaneously to induced recombination. Brains of the animals were collect at postnatal day 4 or day 7 for analyses.

The following primers were used for mouse genotyping: YAP(dupa)-F: AGGACAGCCAGGACTACACAG, YAP(dupa)-R: CACCAGCCTTTAAATTGAGAAC, YAP(hmc)-F: GTCTTTCTCTAGGCACAAAAAGG, YAP(hmc)-R: AGTGGTAAAGAATAATGCTCATCC, TAZ-F: CTTCCAAGGTGCTTCAGAGA, TAZ-R: GGAGAGGTAAAGCCCACCAG, ZsGreen-9103: GGCATTAAAGCAGCGTATCC, ZsGreen-9104: AACCAGAAGTGGCACCTGAC, ZsGreen-9020: AAGGGAGCTGCAGTGGAGTA, ZsGreen-9021: CCGAAAATCTGTGGGAAGTC

All experiments involving animals were conducted under IACUC guidelines at Northwestern University.

### Cell culture

Neural stem cells were dissected at postnatal day 1 mice from the subventricular zone (SVZ) by using forceps to mechanically remove the lateral ganglionic eminence, followed by chemical dissociation using 0.05% trypsin with EDTA. Cells were maintained and passaged in a floating culture with DMEM/F12 media (Gibco/Invitrogen) with N2, B27, and 20ng/mL EGF (Millipore). For differentiation, cells were plated on Poly-D-Lysine (PDL) /Laminin (BD 354087) coated glass coverslips in 0.2ng/mL EGF at a concentration of 50,000 cells per well if not otherwise specified. PDL only glass coverslips (Corning 08-774-384) were used for select experiments that examined the role of β1-integrin signaling. Cells from wild-type mice were used between passage 2 and passage 5.NPCs dissected from transgenic mice were treated with Adeno-Cre (Vector Biolabs 1700) or Adeno-GFP (Vector Biolabs 1060) at a concentration of 1:2000 at 18 hours after passage 2 to ablate the relevant gene of interest. Virus was removed 48 hours after addition. Cells were not used for experiments until at least 7 days after virus addition.

### Cell Treatments

Floating cultures treated with BMP4 (R&D) had 20ng/mL added for 6 hours, after which either protein or RNA was collected. Adherent cultures for staining were treated during plating with 10ng/mL BMP4. For ILK inhibition, cpd22 (Millipore) was administered to floating cultures at 1uM for 6 hours. Adherent cultures were treated during plating with 0.25uM cpd22. MG-132 (Sigma) was added to floating cultures at 100uM to inhibit proteasome-mediated degradation immediately prior to cpd22 treatment, so that cells were treated with both drugs simultaneously. Cycloheximide (CHX) (Sigma) was added to floating cultures at 100ng/mL to inhibit translation immediately prior to BMP4 treatment, so that cells were treated with both drugs simultaneously. β1-integrin blocking antibodies (BD Bisciences 555003) were added to cultures for 24 hours at a concentration of 1:100.

### Nucleofection of NPCs

The plasmid for TAZ(S89A) overexpression (Addgene 19026) was nucleofected using program A-033 using the Lonza Amaxa nucleofector I and the Lonza Mouse NSC kit.

### Western blots

Protein samples were collected in either M-PER (Thermo Fisher 78501) with HALT protease and phosphatase inhibitor (Thermo Fisher 78440) for whole cell lysates, or using a nuclear/cytoplasmic extraction kit (Thermo Scientific 78835) to separate compartments. Protein was boiled in reducing buffer for 20 minutes, then run on SDS-Page gradient gels, transferred to PVDF membrane, blocked for one hour with either 5% Blotto in TBS-Tween or 5% BSA in TBS-Tween, then probed with antibodies in blocking buffer at 4°C overnight. Secondary antibody detection was done on the following morning. Antibodies used for immunoblots included TAZ (BD Pharmigen 560235, 1:500), TAZ (Cell Signaling 83669S, 1:500), pTAZ (CS 59971, 1:500), GFAP (Dako Z0334, 1:1000), GAPDH (Millipore, 1:4000), YAP/TAZ (Sigma WH0010413M1, 1:1000), pSerYAP (CS 4911S, 1:1000), pLATS1 (Cell Signaling, 1:1000), LATS1 (Cell Signaling, 1:1000), pGSK3b (Cell Signaling 9336, 1:500), GSK3b (Cell Signaling 9315, 1:500). Secondary antibodies were used at 1:2000: anti-mouse IgG-HRP (CS 7076), anti-rabbit IgG-HRP (CS 7074). Pierce ECL or Pierce Femto ECL was added for 5 minutes, and blots were imaged either using Amersham Hyperfilm ECL (GE) on the Azure c600 (Azure Biosystems) and quantified using NIH ImageJ software.

### Immunocytochemistry

For *in vitro* studies, cells on glass coverslips were washed with cold 1X PBS, then fixed for 20 minutes at 4°C in 4% paraformaldehyde. Cells were washed three times with 1X PBS, then stored at 4°C until staining. Cells stained for proliferation were fixed at 1 day post-plating, while cells stained for differentiation were fixed at 3 days post-plating. Coverslips were blocked with 10% normal goat serum in PBS with 0.025% Triton-X for one hour at room temperature. For *in vivo* studies, brains tissue fixed in 4% paraformaldehyde were cryoprotected in 30% sucrose for 2 hours followed by embedding in OCT for sectioning to 10μm thickness on a cryostat (Leica CM1860). Prior to staining, antigen retrieval treatment in 10 mM sodium citrate buffer was performed at 95°C for 2 minutes. Sections were then wash 3 times with PBS and blocked with 10% normal goat serum in PBS with 0.25% Triton-X for one hour at room temperature prior to the addition of primary antibodies.

Primary antibodies were diluted in the blocking solution and cell incubated overnight at 4°C. Sections or coverslips were washed 4 times with PBS with 0.025% Triton-X at room temperature after primary incubation. AlexaFluor secondary antibodies were diluted in blocking buffer at 1:1000 with Hoescht nuclear stain at 1:2000, and incubated for one hour at room temperature. After washing 4 times with PBS-Triton-X, sections or coverslips were mounted with Prolong Gold. Primary antibodies used include YAP/TAZ (Sigma WH0010413M1, 1:250), S100β (Proteintech 14146-1-AP, 1:500), GFAP (abcam 4674, 1:1000), Nestin (BD 556309, 1:2000), TAZ (Cell Signaling 83669, 1:500), Aldh1L1 (abcam 87117, 1:500), Olig2 (Millipore Sigma MABN50, 1:500). Cells stained with TAZ were subjected to a 10-minute antigen retrieval treatment in 0.1% SDS at 37°C prior to blocking.

### EdU treatment and staining

For cells, 10μM EdU was administered to cultures four hours before fixation. For *in vivo* labeling, EdU (50mg/kg) was injected on postnatal day 4 four hours before the brains were harvested. Staining was done through use of the Click-iT EdU Cell Proliferation Kit, Alexa Fluor 594 dye (Invitrogen C10339).

### qRT-PCR

RNA was collected from cultured NPCs with the Qiagen RNeasy micro kit (Qiagen 74004) according to the manufacturer’s protocol. cDNA was synthesized using the SuperScript IV kit (Thermo Fischer 18090010), and qPCR conducted using the SYBR Green system (Thermo Fisher 4385612).

Primers used: YAP1-F: CAGAATGCTGCGAGGTCATA, YAP1-R: ATGGCCTCAAATGACTGACC , TAZ-F: GAAGGTGATGAATCAGCCTCTG, TAZ-R: GTTCTGAGTCGGGTGGTTCTG, CTGF-F: GGGCCTCTTCTGCGATTTC, CTGF-R: ATCCAGGCAAGTGCATTGGTA, GAPDH-F: GTC GTG GAT CTG ACG TGC C, GAPDH-R: TGC CTG CTT CAC CAC CTT C.

### Imaging

Images for differentiation experiments were taken on a confocal microscope (Leica SP5) at 40x and exported to NIH Image J for cell counting. Image stacks used for counting contained 4 slices taken 2.5um apart, flattened using max projection. Images for proliferation experiments were taken on a fluorescent microscope (Zeiss Axiovert 200m) at 20x and exported to NIH Image J for cell counting.

### Cell area quantification

Images taken for differentiation experiments were also used for cell area quantification. Images were converted to 16-bit grayscale, threshold for the GFAP channel set to 5, and area in micron^2 automatically measured. Cells touching the border of an image and areas of GFAP-positivity under 10um^2 and over 2000um^2 were excluded in order to maximize the number of cells captured and exclude multiple-cell aggregates.

### Sholl analysis

Images used for differentiation cell counts were also used for Sholl analysis. GFAP+ astrocytic processes semi-manually traced and analyzed following protocols previously established using Fiji (Ferreira et al., 2014). A radius step size of 4um from the center of DAPI-stained nuclei was used.

### Data analysis and statistics

Data analysis and graph generation was conducted using Prism 8 using the statistical tests specified by experiment. When relevant, paired t-tests were used for experiments in which cells treated with Adeno-GFP and Adeno-Cre come from the same individual mouse. Data are all presented as means ± standard error of the mean (s.e.m.).

## Results

### BMP signaling promotes TAZ but not YAP expression in postnatal NPCs

Given that YAP and TAZ have a large amount of sequence homology, and that many antibodies recognize both, we first validated the antibodies we would use for our studies. To accomplish this, we first utilized mutant mice with Cre/LoxP mediated conditional ablation of TAZ and examined TAZ expression in postnatal NPCs. Cultured NPCs from homozygous TAZ mutant (TAZ^fx/fx^) were treated with either an adeno GFP or an adeno Cre virus for 7 days, followed by protein analysis via Western blot using a monoclonal TAZ antibody. We identified a 50 kD TAZ band in the adeno GFP treated control (TAZ^+/+^) cultures that was absent in the TAZ ablated (TAZ^-/-^) cells (Fig. 1A). We then added the adeno Cre virus to cultures of Yap^+/fx^ and YAP^fx/fx^ cells and identified a band at 65kD that was lost in YAP ablated (YAP^-/-^) cultures as YAP (Fig 1B). Another band at 50kD was also detected that was partially reduced in YAP^-/-^ cultures. Since TAZ is detected at 50kD, we asked whether YAP antibody also detects TAZ. Importantly, when probed with a monoclonal TAZ antibody, no reduction in the 50kD band was observed in YAP^-/-^ cells (Extended Data Fig 1-A-C). This finding suggests that the 50kD band detected by the YAP antibody likely represents a different YAP isoform and not TAZ, and that ablation of YAP does not regulate TAZ. To determine if the converse is also true, we probed for YAP in TAZ^-/-^ cells, and found no alteration in protein levels (Extended Data Fig 1-1D-F). This suggests that YAP protein expression is not affected by TAZ ablation in postnatal NPCs.

**Figure 1.**
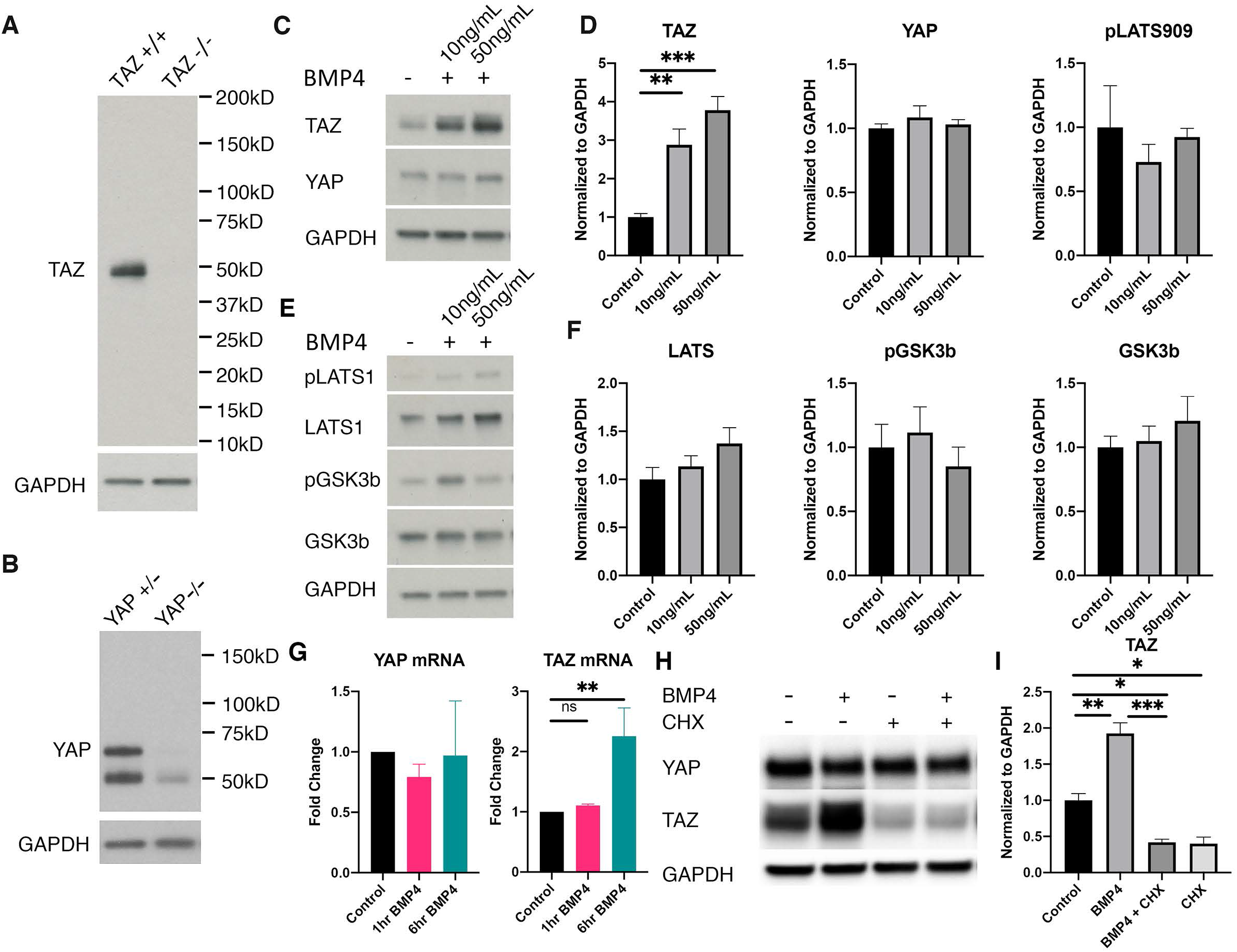
Short-term BMP signaling regulates TAZ, but not YAP, at the post-transcriptional level. A) Validation of TAZ antibody specificity by western blot analysis of TAZ protein expression in TAZ^+/+^ and TAZ^-/-^ NPCs. B) Validation of YAP antibody specificity by western blot analysis of YAP protein expression in YAP^+/+^ and YAP^-/-^ NPCs. C,D) TAZ but not YAP expression in NPCs responds to BMP4 treatment in a dose-dependent manner (TAZ by RM one-way ANOVA, *n*=4, ***p=0.0008; Tukey’s multiple comparisons test: control vs 10ng/mL **p=0.0055, control vs 50ng/mL ***p=0.0007, YAP by RM one-way ANOVA, *n*=4, p=0.3798). E,F) Short-term BMP4 treatment in NPCs does not impact pLATS(S909) (RM one-way ANOVA, p=0.6497), LATS1 (RM one-way ANOVA, p=0.1568), pGSK3b (RM one-way ANOVA, p=0.6016), or GSK3b (RM one-way ANOVA, p=0.4817) protein levels. All *n* = 4. G) BMP (20ng/ml) increases TAZ but not YAP mRNA levels. (TAZ by ordinary one-way ANOVA, **p=0.0076; Dunnett’s multiple comparisons test: control vs 1 hr BMP4 p=0.9088, control vs 6 hrs BMP4 **p=0.0070, YAP by ordinary one-way ANOVA, p=0.7552). All *n*=4. H,I) CHX treatment impairs BMP-mediated TAZ protein increases (RM one-way ANOVA, ****p<0.0001; Tukey’s multiple comparisons test: control vs BMP4 **p=0.0021, control vs BMP4+CHX *p=0.0214, control vs CHX *p=0.0186, BMP4 vs BMP4+CHX ***p=0.0001, BMP4 vs CHX ***p=0.0001, BMP4+CHX vs CHX p=0.9990). All *n* = 3. Data are presented as means ± s.e.m.

Once we validated the specificity of our antibodies, we examined the effects of BMP signaling on TAZ and YAP levels in NPCs. Wildtype (WT) NPC cultures were treated with BMP4 for 6 hours, and YAP and TAZ protein levels were examined via immunoblotting. BMP regulated TAZ protein in a dose-dependent manner, but YAP protein did not respond to treatment (Fig 1C,D). Given that traditional Hippo signaling regulates TAZ and YAP protein stability through LATS phosphorylation, we probed for pLATS1 and total LATS1, but found no significant changes in the relevant time frame (Fig 1D-F), suggesting that BMP-mediated upregulation of TAZ is independent of Hippo signaling. GSK3β, another kinase that regulates TAZ phosphorylation and degradation in other systems (Huang et al., 2012) similarly showed no change (Fig 1E,F) in the presence of BMP. Since YAP and TAZ did not seem to be regulated by traditional LATS phosphorylation and degradation, we next checked mRNA levels by quantitative PCR (qPCR) to see if transcription was affected. No change in YAP mRNA was detected with or without BMP4 treatment (Fig. 1G). By contrast, TAZ mRNA was more abundant after 6 hours of BMP4 treatment, although the increase seems smaller than what would be expected from the observed protein changes (Fig 1G). We therefore examined if translation was instead the more important regulatory step, and found that inhibition of translation through addition of 100ng/mL cycloheximide (CHX) prevented BMP-induced TAZ protein increases. Additionally, CHX alone led to a decrease in TAZ protein, indicating that TAZ has a relatively short half-life. In contrast, no significant change in YAP protein levels was observed after CHX treatment (Fig 1G,I). Taken together, these results indicate that BMP signaling promotes the expression of TAZ, but not YAP, at both mRNA and protein levels in postnatal NPCs.

Since YAP and TAZ must translocate to the nucleus to exert their transcriptional effects, we next examined the effects of BMP signaling on their nuclear presence. A 6-hour BMP4 treatment of NPC cultures resulted in increased levels of TAZ in both the nuclear and the cytoplasmic protein fractions (Fig 2A,B,E,F). No effect of BMP on total YAP was detected in either the cytoplasmic or the nuclear fraction (Fig 2A,C,E,G). We also did not see changes in phosphorylated YAP (Ser127) marked for degradation in the cytoplasm (Fig 2D), nor did we observed altered expression of the YAP/TAZ cofactor TEAD4 in the nucleus (Fig 2H). Immunocytochemical analysis of TAZ protein in adherent NPCs cultured in the presence or absence of BMP4 for 2hrs also demonstrated an increase in TAZ protein expression with BMP4 treatment (Fig. 2I). To examine if the BMP-induced increase in nuclear TAZ was transcriptionally active, we measured mRNA levels of connective tissue growth factor (CTGF), a TAZ transcriptional target and a BMP4 antagonist (Abreu et al., 2002; Zhang et al., 2009), by qPCR in TAZ knockout NPCs with and without BMP4 treatment. CTGF mRNA decreased in TAZ knockout cells, and a BMP4-induced increase in CTGF mRNA CTGF was partially abrogated by TAZ knockout (Fig 2J), suggesting that TAZ mediates some of the effects of BMP signaling on transcriptional targets.

**Figure 2.**
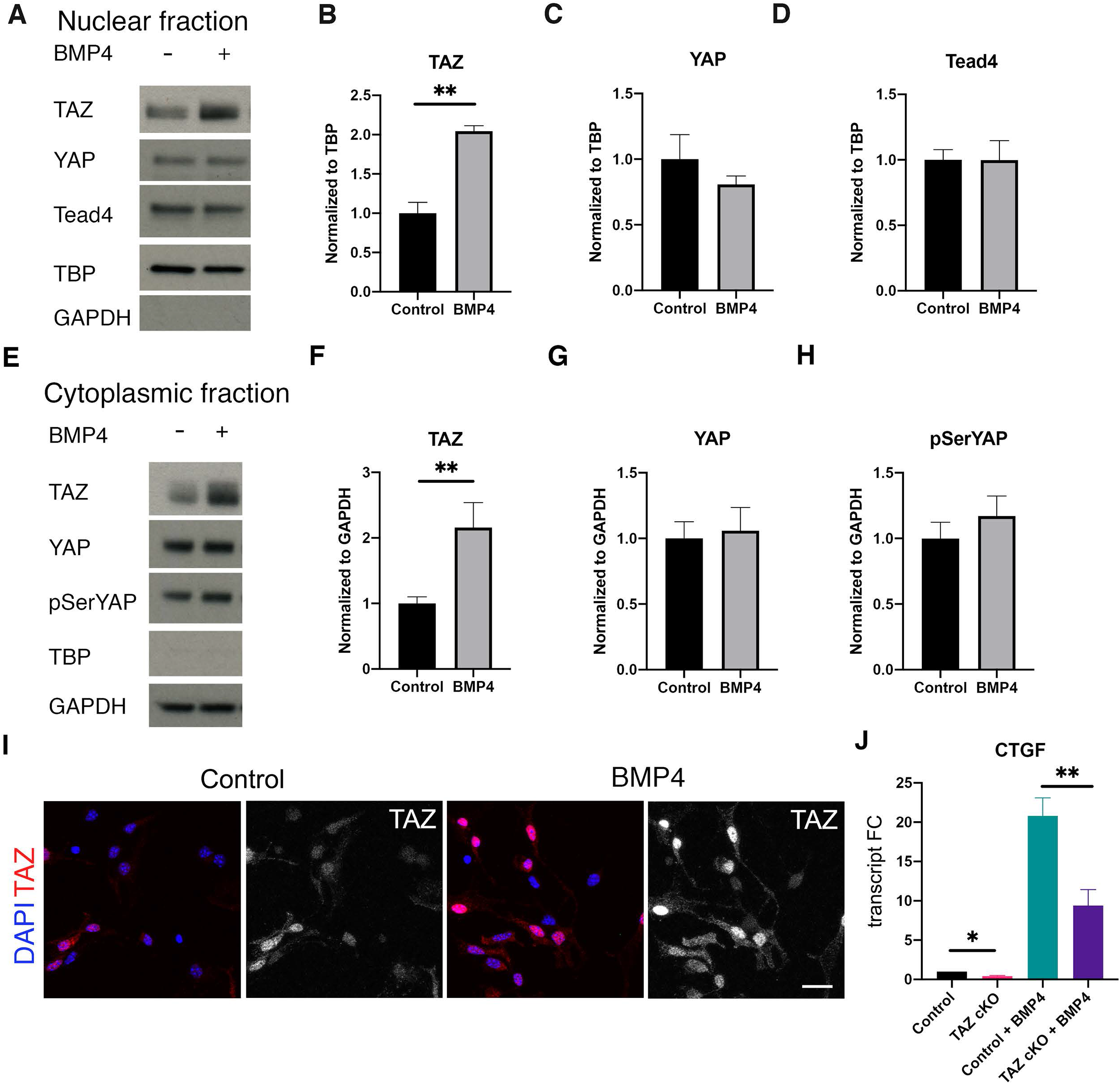
BMP increases TAZ protein levels in both the nuclear and cytoplasmic fractions of NPCs, but does not impact YAP sub cellular localization. A-D) The nuclear fraction of proteins isolated from NPCs treated with BMP4 shows an increase in TAZ, but not YAP (TAZ ratio paired t-test **p=0.0047, YAP ratio paired t-test p=0.2290, TEAD4 ratio paired t-test p=0.7628). All *n=*5. E-H) The cytoplasmic fraction of proteins from BMP4-treated NPCs shows an increase in TAZ, but no change in total YAP or YAP phosphorylated for degradation (TAZ ratio paired t-test **p=0.0058, YAP ratio paired t-test p=0.8858, pSerYAP ratio paired t-test p=0.2263). *n* =6. I) Wild-type NPCs plated on PDL/laminin, treated for 2 hours with 20ng/mL BMP4. A trend towards increased nuclear TAZ can be observed in NPCs plated on PDL/laminin (n=3, paired t-test, p=0.14). Scale bar = 20μm. Data are presented as means±s.e.m. J) CTGF mRNA is increased in response to BMP4 signaling in NPCs, and that response is partially abrogated by deletion of TAZ (by RM one-way ANOVA: F3,6=57.74, ****p<0.0001; paired t-test control vs TAZ cKO *p=0.0260; paired t-test control+BMP4 vs TAZ cKO + BMP4 **p=0.0057). *n*=3.

### TAZ/YAP regulate postnatal astrocytic differentiation in cultured NPC

The observation that BMP signaling increases TAZ levels suggested that TAZ might mediate some of the gliogenic effects of BMP signaling in NPC, such as cell cycle exit and astrocytic differentiation. It was previously reported that Nestin-Cre mediated conditional ablation of YAP in NPCs at embryonic stages led to reduced number of proliferative astrocyte progenitors as well as differentiated astrocytes (Huang et al., 2016b), suggesting that YAP is necessary for the maintenance of astrocyte progenitor pool during embryonic and perinatal development. To investigate the role TAZ and YAP in BMP-mediated postnatal astrocyte differentiation, we first conditionally delete TAZ and YAP in cultured postnatal NPC in the presence and absence of BMP4. TAZ and YAP were both ablated using an adeno-Cre virus in YAP^fx/fx^/TAZ^fx/fx^ cells to generate double conditional knockout (dcKO) NPCs, and EdU incorporation was quantified as a measure of cell division one day post-plating on PDL/laminin coated coverslips in differentiation conditions. Similar to YAP ablation in embryonic NPCs (Huang et al., 2016a), ablation of YAP and TAZ together did not significantly alter EdU incorporation by nestin- expressing postnatal NPCs (Fig 3A,B,I). Since prior studies have posited that YAP mediates the anti-proliferative effects of BMP signaling in NPCs (Yao et al., 2014), we examined the responses of YAP and TAZ dcKO NPCs to BMP4 treatment. Treatment of NPCs with BMP4 for 24 hours in differentiation conditions dramatically reduced EdU incorporation in both control and YAP/TAZ dcKO cultures compared to their non-treated counterparts (Fig 3C,D,J). These findings indicate BMP-induced cell cycle exit takes place independent of TAZ and YAP functions. However, the number of EdU+ cells in BMP4-treated controls was further reduced significantly (p=0.0328, Fig 3J) in YAP/TAZ dcKO cells treated with BMP4, suggesting that YAP and/or TAZ maintain a small population of dividing NPC during BMP induced astrocyte differentiation.

**Figure 3.**
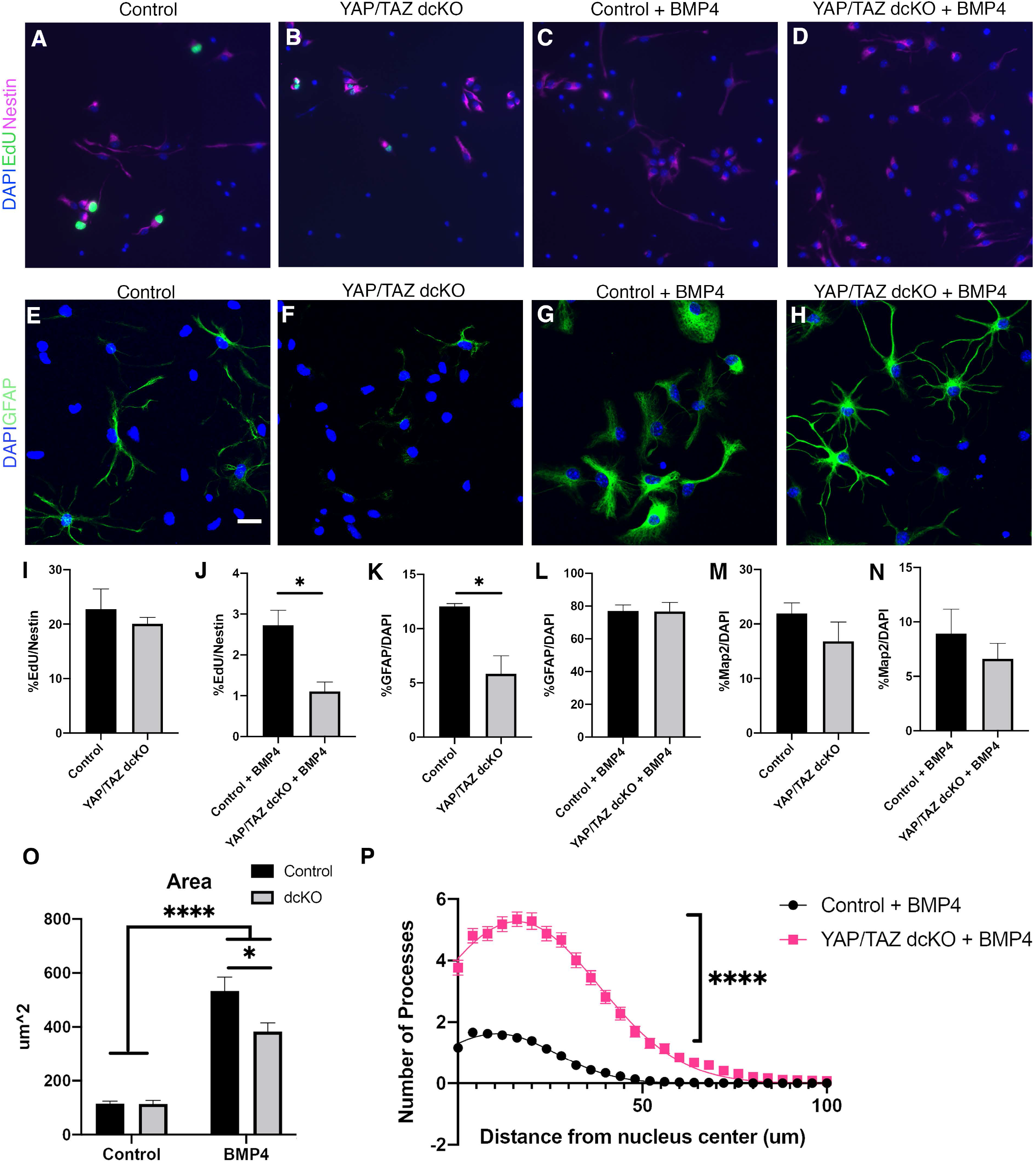
YAP and TAZ do not impact NPC proliferation, but do affect NPC astrocytic differentiation. A,B,I) EdU incorporation in control versus YAP/TAZ double knockout NPCs, plated on PDL (paired t-test p=0.5387, *n*=5). C,D,J) EdU incorporation in control and YAP/TAZ dcKO NPCs treated with 10ng/mL BMP4 for 24 hours on PDL/laminin (paired t-test *p=0.0328, *n*=5). E,F,K) GFAP+ astrocytes differentiated from control versus YAP/TAZ double knockout NPCs after 3DIV on PDL/laminin (paired t-test *p=0.0134 *n*=5). G,H,L) GFAP+ astrocytes generated from BMP4-treated control and YAP/TAZ dcKO cells on PDL/laminin (paired t-test p=0.9595 *n*=5). M) Quantification of MAP2+ cells generated from control and YAP/TAZ double knockout NPCs on PDL/laminin (paired t-test p=0.3881 *n*=5). N) Quantification of MAP2+ cells generated from control and YAP/TAZ double knockout NPCs treated with BMP4 on PDL/laminin (paired t-test p=0.4833 *n* = 5). O) Quantification of cell area for GFAP+ astrocytes. (by two-way ANOVA: BMP4, ****p<0.0001; YAP/TAZ dcKO, p=1.009; BMP4*YAP/TAZ dcKO, p=0.0694; Sidak post-hoc test *YAP/TAZdcKO+BMP4* versus *Control+BMP4* *p<0.05). P) Sholl analysis conducted on control and YAP/TAZ dcKO GFAP+ astrocytes generated in BMP4 on PDL/laminin with a Gaussian fit (unpaired t test of nonlinear fit ****p<0.0001, N=4 or 5, n=20). Data are presented as means ± s.e.m. Scale bars, A-D = 50μm; E-H = 20μm.

We next investigated whether YAP/TAZ affect astrocytic lineage commitment and differentiation. Using GFAP expression as a marker of astrocytic lineage, we found that the number of GFAP+ astrocytes detected after three day *in vitro* differentiation of YAP/TAZ dcKO NPCs is significantly reduced (p=0.0134, Fig 3E,F) without any change in the number of Map2+ cells (p=0.3881, Fig 3K,M) when compared to wild type controls. However, YAP/TAZ dcKO had no effect on the large Increase in GFAP+ astrocytes after treatment with BMP4 (Fig 3L,N), suggesting that YAP/TAZ are not required for the effects of BMP4 on GFAP expression or astrocytic lineage commitment. However, while YAP/TAZ dcKO did not alter the percentage of cells that differentiated into GFAP+ astrocytes after BMP4 treatment, the morphology of the astrocytes generated was strikingly different (Fig 3 G,H). BMP4-induced YAP/TAZ dcKO astrocytes were more stellate and have more elongated processes than BMP4-generated control astrocytes, and the GFAP+ astrocytes in the dcKO condition had a much smaller cytoplasmic volume (Fig 3O). Sholl analysis of GFAP+ cells generated by BMP treatment also showed a significant increase in the number of processes in dcKO compared to control cells (Fig 3P). Interestingly, YAP/TAZ dcKO had no effect on GFAP+ astrocyte numbers or morphology when NPCs were plated on PDL-only coverslips for differentiation (Extended Data Fig 3-1), indicating that the presence of laminin is necessary for TAZ and YAP to impact astrocyte differentiation.

### TAZ is necessary and sufficient for postnatal astrocytic maturation

Since knockout of TAZ and YAP together altered the differentiation of postnatal NPCs, we studied them separately to determine their individual effects on NPC proliferation and astrocytic differentiation. We first examined cell division of postnatal NPCs from TAZ^fx/fx^ mice transduced with either Adeno-GFP virus (control) or Adeno-Cre virus (TAZ cKO). One day post- plating in differentiation conditions, EdU incorporation did not differ significantly between control and TAZ cKO cells (Fig 4A,B,M). Since Hippo pathway components may interact with EGF signaling to exert effects on proliferation (Yang et al., 2012; Fan et al., 2013; Xia et al., 2018), we increased the concentration of EGF in the culture media 10-fold to 10ng/ml to determine whether this would help to elicit effects of TAZ, but there was still no difference between the two groups (data not shown). To examine the gain of function effects of TAZ on NPC proliferation, , we nucleofected NPCs with a constitutively active TAZ, TAZ(S89A), which prevents phosphorylation necessary for TAZ protein degradation and increases nuclear TAZ-mediated transcription (Lei et al., 2008). TAZ(S89A) overexpression did not significantly alter EdU incorporation in NPCs (Fig 4C,D,N) despite an over 12-fold increase in TAZ expression without any accompanying increase in the phosphorylated version (Fig 4S). Thus TAZ had no demonstrable effect on cell cycle state of postnatal NPC.

**Figure 4.**
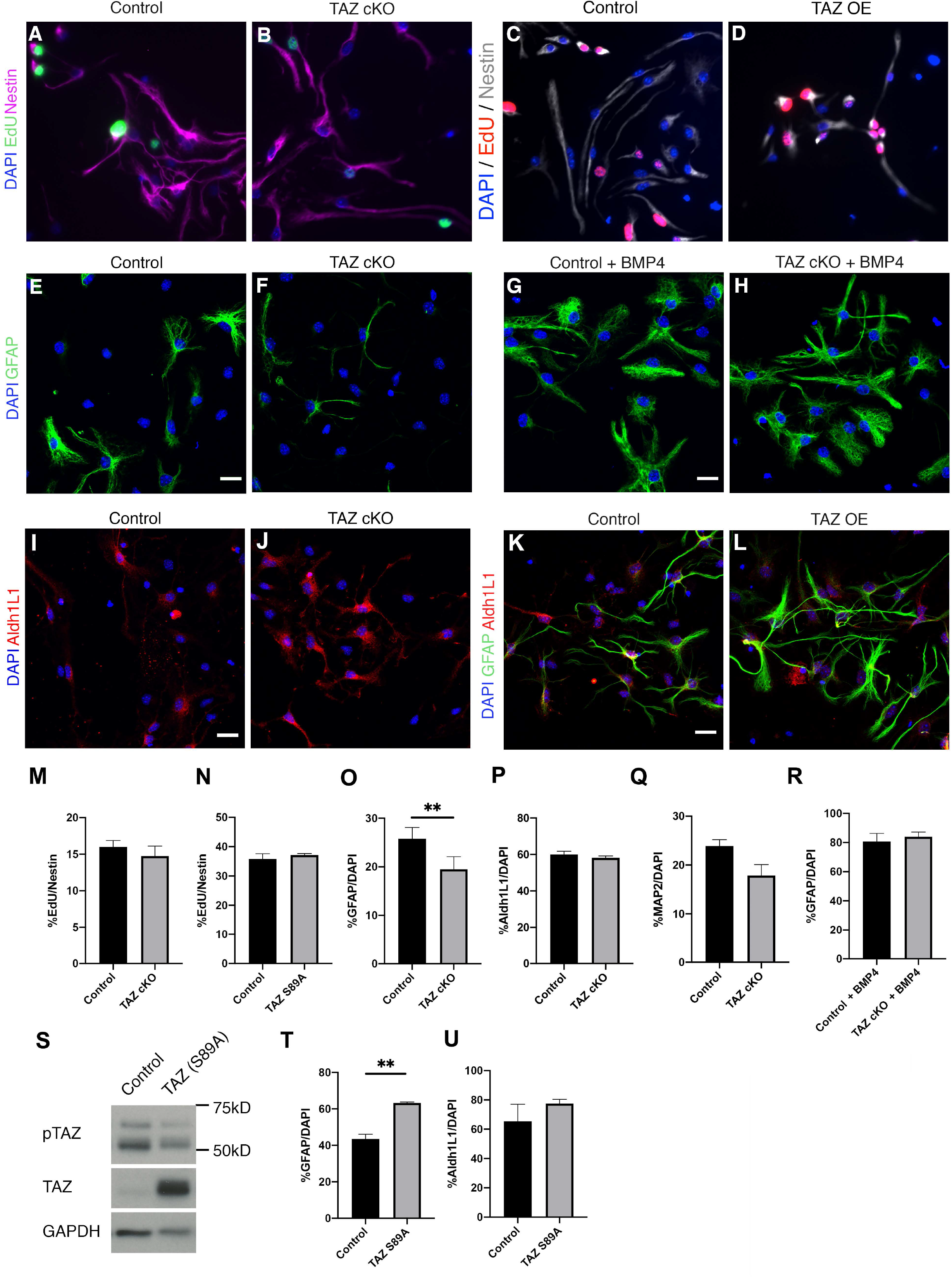
TAZ promotes NPC differentiation into GFAP+ astrocytes. A,B,M) EdU incorporation in control and TAZ knockout cells plated on PDL does not differ (paired t-test p=0.2236 *n* = 4). C,D,J) EdU incorporation in control and TAZ(S89A) overexpression cells plated on PDL does not differ (unpaired t-test p=0.4966 *n* = 3). TAZ knockout cells differentiate into fewer GFAP+ astrocytes (E,F,O, paired t-test **p=0.0078 *n* = 8) without changing the number of Aldh1L1 (I, J, P, paired t-test, p=0.2312, *n* = 8) on PDL/laminin coverslips after 3DIV. G,H,R) GFAP+ astrocytes differentiated from control and TAZ cKO cells in BMP4 treatment (paired t- test p=0.5182 *n* = 8). K,L,T) TAZ overexpression (TAZ OE) results in an increased number of GFAP+ astrocytes after 3DIV (unpaired t-test **p=0.0017 *n* = 3). S) Immunoblot showing overexpression of TAZ(S89A) does not result in increases to the phosphorylated version of TAZ (band around 55kD). U) Aldh1L1 cell counts of control and TAZ overexpressed cells (unpaired t-test p=0.3664 *n* = 3). Data are presented as means ± s.e.m. Scale bars = 20μm.

We next quantify the number of GFAP+ cells to determine the effects of TAZ ablation on astrocyte differentiation. Similar to YAP/TAZ dcKO NPCs, TAZ cKO NPCs generated fewer GFAP+ cells under differentiation conditions when compared to control cells on PDL/laminin (p=0.0078, Fig 4E,F,O), indicating that TAZ alone regulates lineage commitment and/or maturation of astrocytes. To distinguish these possibilities, we quantified the number of cells expressing the pan-astrocytic marker aldehyde dehydrogenase 1L1 (Aldh1L1) and found no significant difference between control and TAZ cKO astrocytes (Fig 4I, J, P), suggesting that TAZ regulates astrocyte maturation and not astrocytic lineage commitment. We also did not observe changes in MAP2 expression, consistent with the notion that neural cell lineages are not affected by TAZ ablation (Fig 4Q). Also similar to YAP/TAZ dcKO NPCs, the number of GFAP- positive cells did not differ between BMP-treated control and BMP-treated TAZ cKO cultures (Fig 4G,H,R). Unlike YAP/TAZ dcKO cells, GFAP+ astrocyte morphology did not significantly differ in the absence of TAZ when cells were differentiated with BMP (Fig 4G,H). However, we found that overexpression of TAZ(S89A) significantly increased the percentage of GFAP+ cells after 3 days in culture (p=0.0017, Fig 4K,L,T). Interestingly, the number Aldh1L1 did not change with TAZ(S89A) overexpression (Fig 4K,L,U), suggesting that gain of TAZ function does not alter astrocyte lineage commitment. These findings suggest that TAZ specifically regulates either GFAP expression or cellular maturation, and not proliferation or morphology of postnatal NPC- derived astrocytes.

By contrast, adeno-Cre virus mediated ablation of YAP (YAP cKO) reduced proliferation of postnatal NPC but did not significantly alter the number of GFAP+ cells (Extended Data Fig 4-1). Further, control and YAP cKO NPCs showed no difference in the number of GFAP+ astrocytes after 3 days of BMP4 treatment, nor did they show any obvious differences in morphology. We also observed no difference in the number of cells expressing Aldh1L1, between control and YAP cKO conditions. Thus in contradistinction to prior reports on the role of YAP in embryonic NPCs (Huang et al., 2016a), ablation of YAP in postnatal NPCs does not affect astrocytic lineage commitment and appears to exert little effect on astrocytic differentiation.

### TAZ and YAP differentially regulate glial progenitors in developing mouse neocortex

Astrogliogenesis in the developing mammalian cerebral cortex begins with lineage commitment of radial glial progenitors in the subventricular zone in late embryogenesis and continues into early postnatal days with local divisions and maturation of astrocyte progenitors in the cortical parenchyma (Kriegstein and Alvarez-Buylla 2009, Molofsky and Deneen 2015, Clavreul et al., 2022) . To investigate how TAZ and YAP regulate postnatal glial progenitor cell fate and astrocyte differentiation, we induced Cre-recombinase mediated deletion of TAZ or YAP/TAZ in Nestin-expressing cells at postnatal day 1 and examined zsGreen reporter-marked recombined cells for changes in astrocytic differentiation in the dorsal neocortex at postnatal day 4. Examination of the number and distribution of zsGreen-expressing cells in the neocortex, where significant number of Nestin-expressing NPC undergo local divisions to generate restricted astrocytic progenitors and postmitotic astrocytes, did not reveal significant differences in the percentage of astrocytic lineage committed GFAP^+^zsGreen^+^ or Aldh1L1^+^zsGreen^+^ cells in TAZ cKO or YAP/TAZ dcKO when compared to TAZ^+/-^ littermate controls (Extended Data Fig 5-1A-C, G). Since TAZ and YAP are known to regulate progenitor proliferation, we next analyzed whether deletion of TAZ or YAP/TAZ alter the number of dividing NPCs. Quantification of proliferating cell nuclear antigen (PCNA)-expressing cells in recombined zsGreen^+^ cells and Aldh1L1^+^zsGreen^+^ astrocytic cells in cortical layers 2-6 revealed no significant change in the percentage of zsGreen^+^ and Aldh1L1^+^zsGreen^+^ cells expressing PCNA (Extended Fig 5-1D-F, H), suggesting that early postnatal ablation of TAZ or YAP/TAZ in NPC did not affect their cell cycle states.

To examine the roles of TAZ and YAP in BMP signaling mediated astrocytic differentiation and maturation, we focused on astrocytic lineage cells in cortical layer 1, where robust nuclear pSMAD1/5/8 immunostaining indicated active BMP signaling (Extended Data Fig 5-1I, I’) due to its proximity to meningeal fibroblasts (Choe et al., 2012; DeSisto et al., 2020). we found that in TAZ cKO mice, there was a near significant increase in the percentage of zsGreen^+^ cells expressing the astrocytic lineage marker Aldh1L1 over TAZ^+/-^ littermate controls (Fig 5A,B, p = 0.0513). However, no appreciable change in the number of Aldh1L1^+^zsGreen^+^ cells was observed in YAP/TAZ dcKO mice compared to the controls (Fig 5A,B, p= 0.707), suggesting that YAP deletion inhibit the increase of Aldh1L1+ cells observed in TAZ cKO mice.

**Figure 5.**
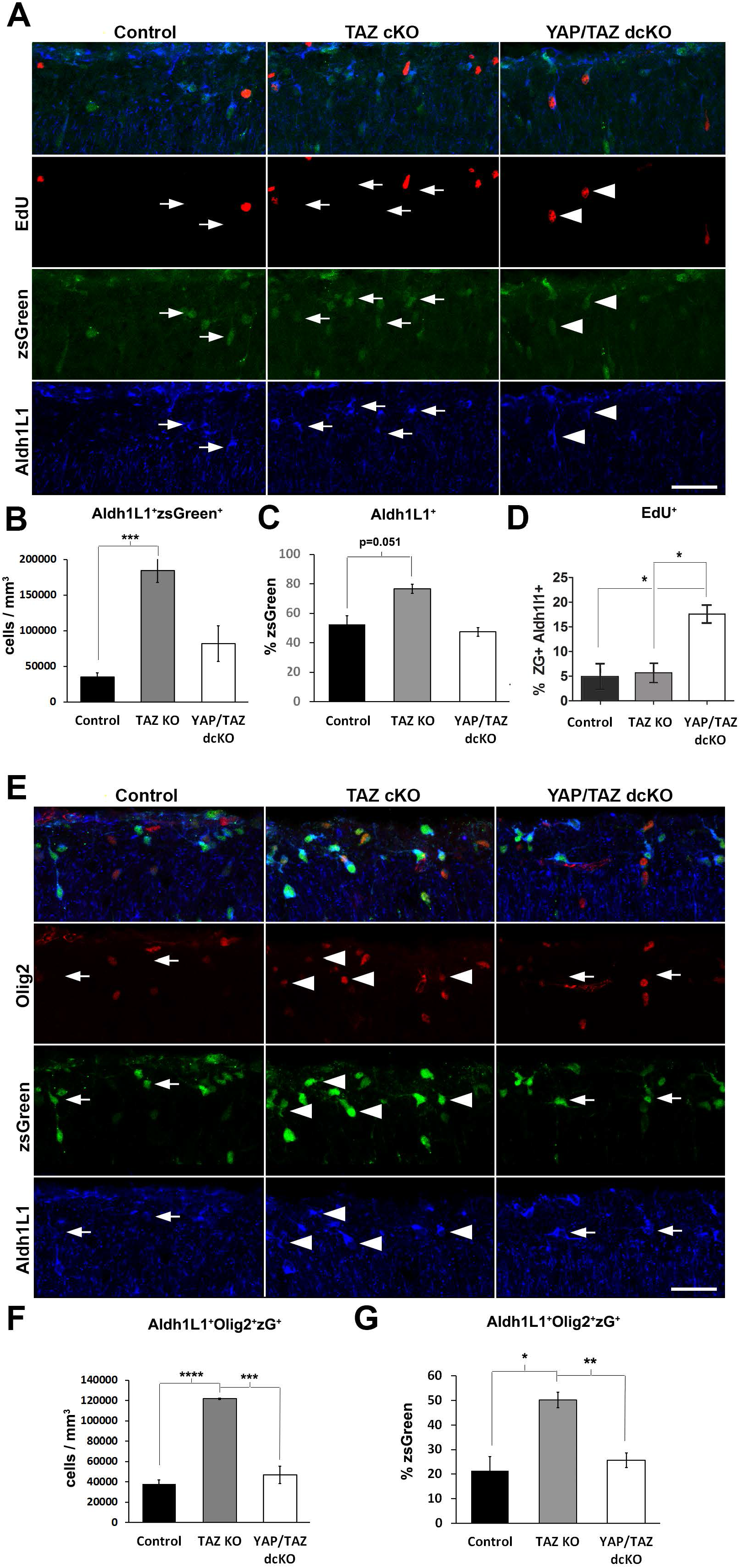
TAZ regulates astrocytic fate specification, not proliferation of intermediate progenitors in developing cortex. A-C) Representative images of cortical layer 1 region in postnatal day 4 mouse brains from control (TAZ^+/-^, *n*=4), TAZ cKO (*n*=3), and YAP/TAZ dcKO (*n*=3) mice stained with astrocyte lineage marker Aldh1L1 (A, blue) and cell cycle marker EdU (A, red) showing an increase in the total number of Aldh1L1^+^zsGreen^+^ cells (A, green) in TAZ cKO when compared with controls (B, one-way ANOVA: F_2,7_, Bonferroni-Holm posthoc, *** p=0.0004). No significant difference in Aldh1L1^+^zsGreen^+^ cells was detected in YAP/TAZ dcKO compared to controls (p=0.122). A similar but not statistically significant increase over controls was also observed in TAZ cKO mice in the percentage of zsGreen+ cells expressing Aldh1L1 (C, one-way ANOVA, F_2,7_, Bonferroni-Holm posthoc, p=0.051). Quantification of the percentage of EdU^+^ cells in Aldh1L1^+^zsGreen^+^ (zG^+^) cells revealed significant higher numbers in YAP/TAZ dcKO when compared to both controls (D, one-way ANOVA, F_2,6_, *p=0.016) and TAZ cKO (one- way ANOVA, *p=0.011). (E-G) Representative images and quantification of Olig2 (D, red) and Aldh1L1 (D, blue) expressing cells in postnatal day 4 neocortex of control (*n*=4), TAZ (*n*=3), and YAP/TAZ deleted (*n*=3) mice. The total number Aldh1L1^+^Olig2^+^zsGreen+ cells was significantly increased in cortical layer 1 of TAZ cKO over controls (F, one-way ANOVA, F_2,7_, Bonferroni-Holm posthoc, **** p=1.06E-05) and YAP/TAZ dcKO mice (*** p=0.00166). Similar changes were detected when Aldh1L1+Olig2+ cells were quantified as a percentage of recombined zsGreen+ cells when compared TAZ ckO with control (G, 1 way ANOVA, F_2,7_, Bonferroni-Holm posthoc, *p=0.0116), or YAP/TAZ dcKO mice (G, 1 way ANOVA, F_2,7_, Bonferroni-Holm posthoc, **p=0.0081). Data are presented as means ± s.e.m. Scale bars = 50μm

We next examined the number of dividing progenitors present in layer 1 across genotypes via quantification of cells incorporating EdU that was injected 4 hours before tissue collection. Despite detecting an increase in Aldh1L1^+^ cells in TAZ cKO, we found no significant changes in the percentage of Aldh1L1^+^ cells expressing EdU (p=0.83, Fig 5A,C). However, a significant increase in the percentage of EdU-expressing Aldh1L1^+^ cells in YAP/TAZ dcKO was observed over controls (p=0.0161) and TAZ cKO (p=0.025) (Fig 5A,C). The difference in EdU^+^Aldh1L1^+^ cells between TAZ cKO and YAP/TAZ dcKO suggests that YAP deletion promoted cell divisions of Aldh1L1+ astrocyte progenitors. Taken together, these findings suggest TAZ and YAP respectively regulate astrocytic differentiation and cell division of BMP signaling-responsive local astrocyte progenitors in cortical layer 1.

To better understand astrocytic lineage progression and maturation from astrocyte progenitors to post-mitotic astrocytes, we next performed co-labeling of Aldh1L1 with the glial progenitor marker Olig2, which has been shown to be transiently expressed in early postnatal astrocyte during differentiation (Cai et al., 2007; Zu et al., 2012). Quantification of zsGreen+ cells that expressed Aldh1L1 and Olig2 showed that TAZ cKO have significantly more zsGreen^+^ recombined cells that co-express Olig2 and Aldh1L1 in cortical layer 1 over controls (Fig 5E-G), both in total number (TAZ cKO vs. control, p=1.06E-5, TAZ ckO vs YAP/TAZ dcKO p=0.00166, Fig 5F) and as a percentage of recombined zsGreen+ cells (TAZ cKO vs. control, p=0.0116, TAZ ckO vs YAP/TAZ dcKO p=0.0081, Fig 5G) TA). These finding suggest that loss of TAZ may either cause precocious generation of Olig2^+^Aldh1L1^+^ astrocyte progenitors or a failure in terminal astrocytic differentiation. *In toto,* these findings show that TAZ, but not YAP deletion in the developing neocortex impairs commitment of glial progenitor to differentiated astrocytes without affecting proliferation.

### Integrin signaling regulates TAZ and YAP expression in postnatal NPCs

β1-integrin promotes stem cell survival and regulates NPC proliferation and lineage commitment (Leone et al., 2005; Pan et al., 2014; Brooker et al., 2016). In other systems, β1- integrin and its downstream signaling components regulate TAZ and YAP through suppression of the Hippo pathway. For example,β1-integrin regulates YAP in mammary epithelial cells (Lee et al., 2019), myofibroblasts (Martin et al., 2016), osteoblasts, and mouse embryonic fibroblasts (Sabra et al., 2017). Downstream effectors of integrins also regulate TAZ nuclear localization (Xia et al., 2019), but extracellular matrix (ECM) effects on TAZ have not been as well examined as those of its paralogue YAP.

To examine effects of β1-integrin signaling on TAZ and YAP expression, we added a β1-integrin blocking antibody to NPC cultures for 24 hours before harvesting the cells for western blot analyses. TAZ protein levels were significantly reduced by treatment with the blocking antibody, whereas YAP and its phosphorylated version were not significantly altered (Fig 6A,C,D,E). TAZ and YAP expression are regulated by a number of kinases including LATS1 and GSK3β, but levels of phospho-LATS1 (Fig 6A,B), phospho-GSK3β, and total GSK3β were not altered (Extended Data Fig 6-1G-I). Similar effects were seen when cells were treated for 6 hours with cpd22, a pharmacologic inhibitor that targets integrin-linked kinase (ILK), a downstream effector of β1-integrin (Extended Data Fig 6-1J-L). To determine if the cellular localization of TAZ or YAP is affected by disruption of β1-integrin signaling, we next fractionated cells and examined the nuclear and cytoplasmic compartments separately. Both nuclear and cytoplasmic fractions of TAZ were significantly decreased in cpd22-treated NPCs. However, only nuclear YAP was decreased in the presence of cpd22 (Fig 6F-H,J-L) suggesting that ILK signaling regulates nuclear shuttling of transcriptionally active YAP.

**Figure 6.**
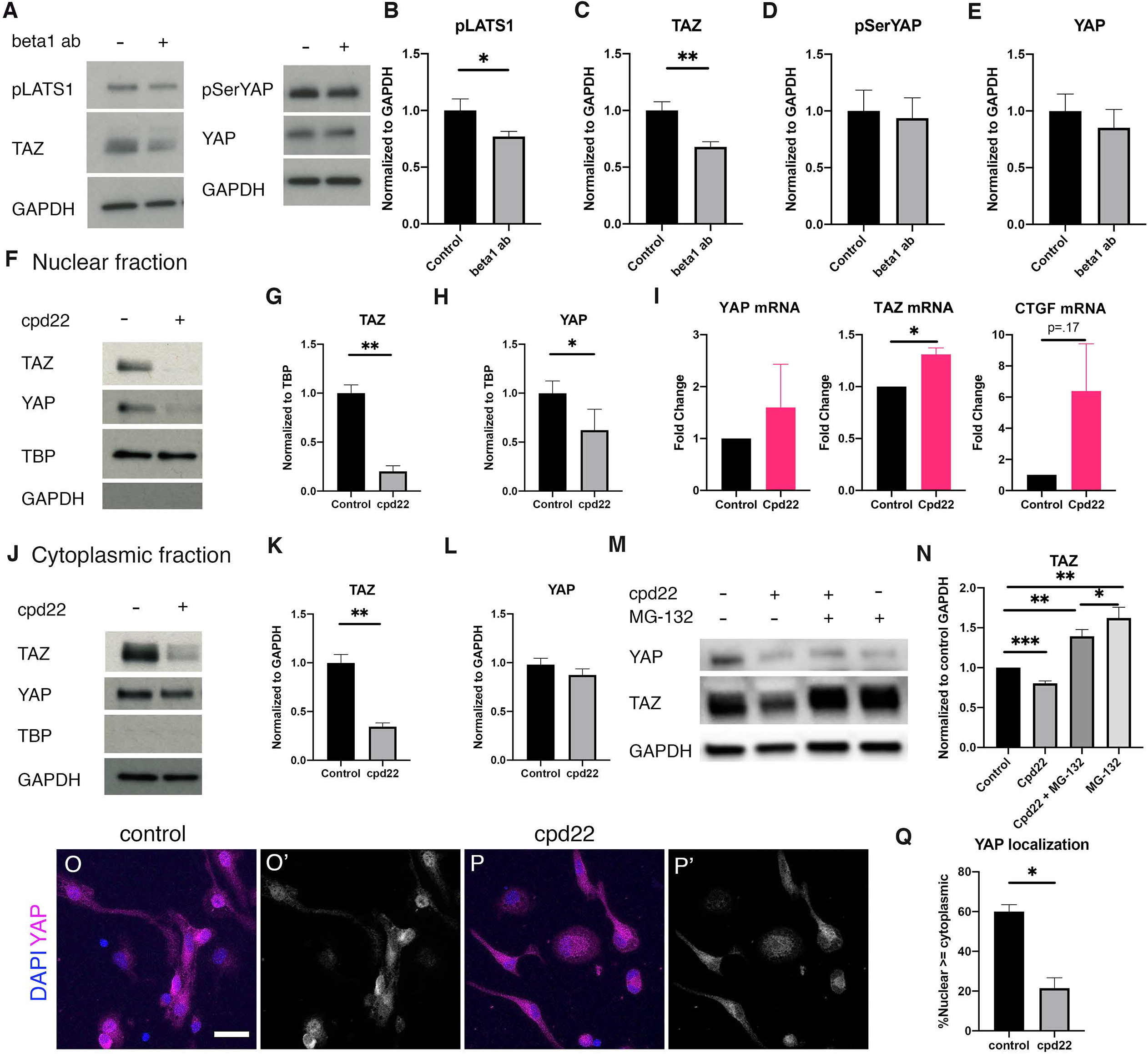
ILK signaling promotes YAP and TAZ protein stability. A-E) 24 hour addition of β1-integrin blocking antibody to NPCs measured by immunoblotting. B) pLATS1 protein levels (paired t-test *p=0.0299 *n*=5). C) TAZ protein levels are reduced by β1-integrin blocking antibodies (paired t-test **p=0.0057 *n*=5). D) pSerYAP protein levels (paired t-test p=0.5974 *n*=5). E) YAP protein levels (paired t-test p=0.1108). F) 6 hour treatment with cpd22 reduces YAP and TAZ protein levels in the nuclear fraction of NPCs. G) TAZ nuclear protein levels (paired t-test **p=0.0021 *n*=5). H) YAP nuclear protein levels (paired t-test *p=0.0409 *n*=5). I) mRNA levels of YAP, TAZ, and CTGF in response to 6 hours of cpd22 treatment (paired t-test; YAP p=0.5251, TAZ *p=0.0163, CTGF p=0.1742. All *n*=4). J) immunoblotting of cytoplasmic fraction of NPCs treated with cpd22 for 6 hours. K) TAZ cytoplasmic protein levels (paired t- test **p=0.0028 *n*=5). L) YAP cytoplasmic protein levels (paired t-test p=0.1047 *n*=5). M) NPCs treated with cpd22, MG-132, or both for 6 hours and probed for TAZ and YAP via immunoblotting. N) TAZ protein levels (via paired t-test; control vs cpd22 ***p=0.0005, control vs cpd22+MG-132 **0.0034, control vs MG-132 **p=0.0036, MG-132 vs cpd22+MG-132 *p=0.0378, *n*=7). O-O’) YAP staining of wild-type NPCs plated on PDL in 2ng/mL EGF. Arrows indicate nuclear localization of YAP in individual cells. Scale bar = 20μm. P-P’) YAP staining of wild-type NPCs treated with 1uM cpd22 for 4 hours on PDL in 2ng/mL EGF. Arrows indicate cells with lower nuclear YAP compared to cytoplasmic YAP. Q) YAP demonstrates decreased nuclear localization in cpd22-treated NPCs after 4 hours (paired t-test *p=0.027, *n*=3). Data are presented as means ± s.e.m.

We sought to examine if transcriptional changes were responsible for the decrease in protein levels. However, cpd22 treatment did not significantly change YAP mRNA, and slightly increased levels of TAZ mRNA (Fig 6I). Staining of cells using a YAP antibody also confirmed that cpd22 treatment for 4 hours was sufficient to reduce YAP levels in the nuclear compartment compared to the cytoplasm (Fig 6O-Q). To determine if the cpd22-mediated decrease in TAZ reflected altered protein turnover, we treated NPCs with MG-132, a proteasome inhibitor, and found that MG-132 treatment abrogated cpd22-mediated degradation of TAZ, but not YAP protein (Fig 6M,N). Together, these data indicate that ILK signaling promotes TAZ protein stability but does not alter nuclear vs cytoplasmic localization of TAZ.

### Integrin signaling promotes NPC proliferation through YAP and inhibits astrocytic differentiation through TAZ

We next asked whether β1-integrin-mediated changes in nuclear/cytoplasmic shuttling of YAP have functional consequences on NPC proliferation or fate specification. Since laminin is a ligand for β1-integrin (Colognato et al., 1997; Belkin and Stepp, 2000; Chernousov et al., 2007), we compared NPCs plated for 24hrs on PDL versus PDL/laminin. As expected, we found that YAP nuclear levels in cells plated on PDL/laminin were slightly more elevated compared to cytoplasmic levels (Fig 7A,C). Additionally, the percentage of Ki67+ cells was increased in the PDL/laminin condition, indicating an increase of cells in cell cycle, consistent with β1-integrin’s role in maintaining stemness of NPCs (Fig 7B) (Campos et al., 2004; Leone et al., 2005). Strikingly, when YAP knockout NPCs were plated on PDL/laminin coated coverslips, they demonstrated reduced EdU incorporation compared to control cells (Fig 7D). This difference in proliferation was not observed when we compared control cells to YAP cKO cells plated on PDL (Fig 7E), suggesting that activation of extracellular matrix-sensing cellular components may be necessary for YAP action, in line with the known roles of YAP and TAZ in mechanosensing. Ablation of TAZ on PDL/laminin did not demonstrate a similar effect on proliferation (Fig 7F), indicating this is a YAP-specific effect.

**Figure 7.**
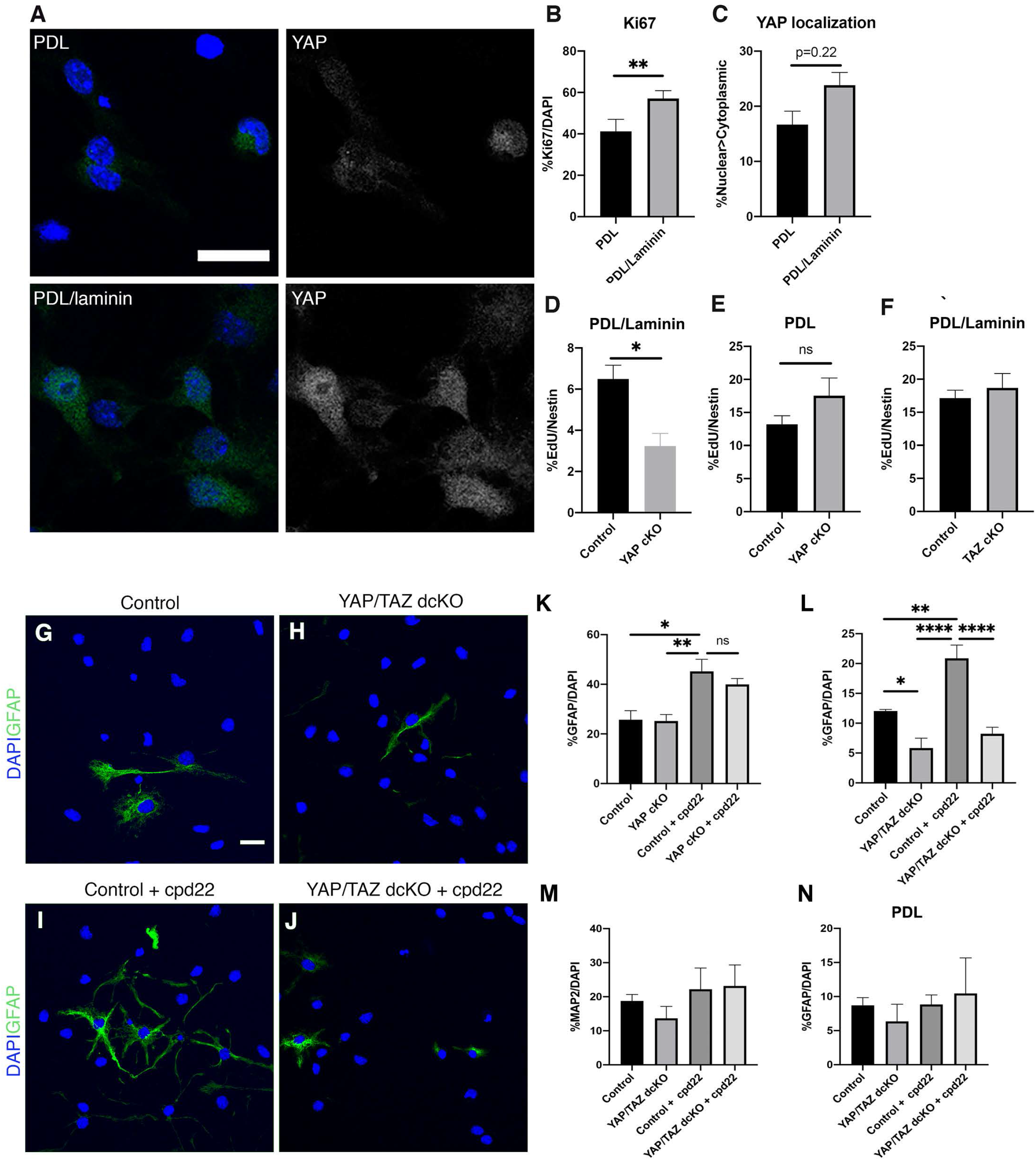
β1-integrin signaling promotes YAP to regulate NPC proliferation and TAZ for astrocytic differentiation. A) NPCs plated on PDL and PDL/laminin stained for YAP. Scale bar = 20μm. B) Ki67+ cells were increased in PDL/laminin plated NPCs compared to PDL-only cells (paired t-test **p=0.0088 *n*=4). C) YAP localizes more to the nucleus of PDL/laminin NPCs compared to PDL-only plated NPCs (paired t-test p=0.2202 *n*=4). D) EdU incorporation is reduced in YAP cKO NPCs plated on PDL/laminin compared to control NPCs (paired t-test *p=0.0406 *n*=5). E) EdU incorporation in control and YAP cKO NPCs on PDL does not differ (paired t-test p=0.0908 *n* = 5). F) EdU incorporation does not differ between control and TAZ cKO NPCs plated on PDL/laminin (paired t-test p=0.5088 *n*=5). G-J) GFAP+ cells generated from control and YAP/TAZ dcKO cells differentiated on PDL/laminin in the presence of cpd22. Scale bar = 20μm. K) quantification of GFAP+ cells in G-J (RM one-way ANOVA F_3,12_=25.42 ****p<0.0001; Tukey’s post-hoc test control vs YAP/TAZ dcKO *p=0.0256, control vs control+cpd22 **p=0.0022, YAP/TAZ dcKO vs control+cpd22 ****p<0.0001, control+cpd22 vs YAP/TAZ dcKO+cpd22 ****p<0.0001, *n*=5). L) quantification of GFAP+ cells generated from cpd22-treated control and YAP cKO cells on PDL/laminin (RM one-way ANOVA F_3,12_=7.786 **p=0.0038; Tukey’s post-hoc test control vs control+cpd22 *p=0.0117, YAP cKO vs control+cpd22 **p=0.0099, *n*=5). M) quantification of MAP2+ neurons in control and YAP/TAZ dcKO NPCs differentiated for 3 days on PDL/laminin with and without ILK inhibitor cpd22 (by ordinary one-way ANOVA p=0.5127). N) quantification of GFAP+ astrocytes in control and YAP/TAZ dcKO NPCs differentiated for 3 days on PDL with and without ILK inhibitor cpd22 (by RM one-way ANOVA p=0.7793). Data are presented as means ± s.e.m.

We also examined interactions between β1-integrin signaling and YAP/TAZ during NPC astrocytic differentiation. Treatment of NPCs on PDL/laminin with cpd22 to abrogate ILK signaling increased the number of GFAP+ astrocytes when compared to untreated controls (Fig 7G, I, L, K), consistent with the known role of β-integrin and ILK signaling in suppressing astrocyte differentiation (Brooker et al., 2016). Knockout of YAP did not alter the effects of cpd22 (Fig 7K). However, GFAP+ astrocyte numbers did not increase after cpd22 in YAP/TAZ dcKO NPCs (Fig 7L). Neuronal differentiation marked by MAP2+ cells was not affected in YAP/TAZ dcKO NPCs (Fig 7M), and the difference in GFAP expression in response to cpd22 was not seen in NPCs plated on PDL alone (Fig 7N). Thus the presence of extracellular laminin promotes YAP/TAZ-mediated inhibition of astrocyte differentiation.

## Discussion

YAP and TAZ regulate stem cell proliferation and lineage commitment in multiple systems, but only until recently have YAP and TAZ been examined separately in the same system. TAZ in particular has not previously been well characterized in neural stem cells. In this study, we examined the roles of YAP and TAZ in astrocytic lineage development, both independently and in the context of BMP and β1-integrin signaling pathways. We found that BMP and Integrin signaling differentially regulate TAZ and YAP expression in NPCs, and that postnatal ablation of TAZ and YAP led to distinct cellular phenotypes in the context of neural progenitor proliferation and astrocyte maturation both *in vitro* and *in vivo*. These findings underline the need to treat these two transcriptional coactivators as distinct molecular entities in the regulation of astrocyte differentiation.

Prior studies on the function of YAP in neural progenitors have revealed roles in both maintenance of stem cell proliferation (Yao et al., 2014) and astrocyte differentiation (Huang et al., 2016). Several key factors differentiate this study from the prior reports. First, both prior reports examined NPCs isolated from day 14.5 embryo (e14.5), whereas this study focused on postnatal day 1 (P1) NPCs. The key difference between e14.5 and P1 NPCs is their lineage potential, as NPCs are transitioning from neurogenic to gliogenic at e14.5 and are only gliogenic at P1. This developmental difference is accompanied by significant transcriptomic and epigenetic level changes in the NPC, which may explain some of the discrepancies between this and prior studies.

Second, Huang et al. utilized Nestin^Cre^ mice (JAX#0037711) to conditionally ablate YAP in NPCs, resulting in YAP ablation as early as embryonic day 11 when Nestin expression is detected in the developing neural tube according to the strain description. Since YAP was shown to maintain NPC proliferation, this early stage YAP deletion likely led to reduced number of late embryonic NPCs, contributing to the observed reduction in gliogenesis at postnatal stages. In contrast, this study utilized a Tamoxifen-inducible Nestin^CreERT2^ mouse line for cell type- and temporal-specific ablation of YAP and TAZ *in vivo*, which yielded some divergent findings when compared to prior reports. Specifically, we found neither YAP or TAZ are necessary for proliferation or differentiation of glial-lineage committed progenitors in layers 2-6 dorsal cortex although astrocyte morphology was altered in the joint absence of TAZ and YAP. This is consistent with past studies on the effect of TAZ and YAP on regulation of cell morphology through cytoskeletal elements, and the lack of striking morphological changes observed with deletion of either one alone suggests that they have overlapping functions in this regard. This also stands in contrast to previous studies that report BMP regulation of YAP activity through phosphorylation (Huang et al., 2016). Third, although less likely, it is also possible that differences between this and prior studies reflects our use of BMP4 rather than BMP2 that was used in the prior studies and/or differences in dosing and time points.

Both our *in vitro* and *in vivo* analyses suggest that ablation of TAZ but not YAP inhibits astrocyte maturation, and that overexpression of TAZ *in vitro* conversely is sufficient to promote astrocyte maturation. However, a significant divergent finding between *in vitro* and *in vivo* studies is the number of Aldh1L1^+^ glial lineage committed cells, which was increased *in vivo* and unchanged *in vitro*. The increase of Aldh1L1^+^ cells observed *in vivo* is in part attributed to a significant increase in Olig2-expressing glial progenitors, suggesting that intermediate progenitors fail to terminally differentiate in the absence of TAZ in the developing brain. The lack of increase in Aldh1L1^+^ cells *in vitro* is likely due to the culture conditions used in our differentiation experiments. This hypothesis is supported by our observation that the effects of TAZ on astrocyte differentiation were only observed in NPCs plated on PDL/laminin-coated coverslips for differentiation, rather than PDL-only coverslips. Combined with our results on EdU incorporation and astrocyte differentiation in YAP/TAZ deleted cells in the presence of ILK inhibitor cpd22, this indicates that YAP and TAZ only have effects on cell proliferation and astrocyte subtype specification in the presence of extracellular matrix activated signals such as β-integrin and ILK. This highlights the importance of carefully controlling culture conditions when examining these signaling pathways *in vitro*.

Our ability to separate YAP and TAZ in this study has enabled us identify that YAP and TAZ are distinct entities with unique roles in NPC fate determination. When downstream of integrin signaling, YAP promotes proliferation, whereas TAZ promotes GFAP-expressing astrocytes. In addition, regulation of these transcriptional coactivators differs. While both are downstream of β-integrin/ILK signaling, BMP signaling regulates TAZ but not YAP, contradicting previous studies. Nevertheless, although BMP signaling targets TAZ into the nucleus with the potential to subserve an instructive role for transcriptional activation and fate determination, TAZ is not required for BMP-mediated promotion of astrocyte lineage commitment. However, BMP signaling has also been shown to regulate maturation of developing astrocyte (Scholtz et al., 2014). Our findings revealed that TAZ modulates GFAP expression post differentiation, suggesting that it could mediate other BMP-regulated maturation properties. CTGF, a BMP regulated gene and known transcriptional target of TAZ, has been shown to be elevated in reactive astrocytes post brain injury and promotes GFAP expression and inflammatory responses(Schwab et al., 2000; Lu et al., 2019). Future studies examining how extracellular mechanosensory cues modify cellular structure through TAZ signaling may provide insights into astrocytic properties during development and post injury.

## Author contributions

JC, YHT, AKL, and CP conducted experiments for the paper. JC, JAK, and CP designed the project and experiments. JC and CP wrote the paper with edits from YHT, AKL, JAK.

## Data Availability Statement

The data that support the findings of this study are available from the corresponding author upon reasonable request.

## Acknowledgments

We would like to thank Mary and Mark Nagelvoort for their generous support.

